# Evidence for the rhythmic perceptual sampling of auditory scenes

**DOI:** 10.1101/618652

**Authors:** Christoph Kayser

## Abstract

Converging results suggest that perception is controlled by rhythmic processes in the brain. In the auditory domain, neuroimaging studies show that the perception of brief sounds is shaped by rhythmic activity prior to the stimulus and electrophysiological recordings have linked delta band (1-2 Hz) activity to the functioning of individual neurons. These results have promoted theories of rhythmic modes of listening and generally suggest that the perceptually relevant encoding of acoustic information is structured by rhythmic processes along auditory pathways. A prediction from this perspective – which so far has not been tested – is that such rhythmic processes also shape how acoustic information is combined over time to judge extended soundscapes. The present study was designed to directly test this prediction. Human participants judged the overall change in perceived frequency content in temporally extended (1.2 to 1.8 s) soundscapes, while the perceptual use of the available sensory evidence was quantified using psychophysical reverse correlation. Model-based analysis of individual participant’s perceptual weights revealed a rich temporal structure, including linear trends, a U-shaped profile tied to the overall stimulus duration, and importantly, rhythmic components at the time scale of 1 to 2Hz. The collective evidence found here across four versions of the experiment supports the notion that rhythmic processes operating on the delta band time scale structure how perception samples temporally extended acoustic scenes.

## Introduction

Perception seems to be systematically controlled by rhythmic processes in the brain [1, 2]. These rhythmic processes may for example reflect the excitability sensory neurons [3-5], the selection of specific features for a behavioral response [6, 7], or the attentional modulation of perception [8, 9]. The perceptually-relevant rhythmic brain activity not only manifests in systematic relations between brain and behavior, such as better perceptual detection rates following a specific pattern of brain activity [10-12], but can also reflect directly in behavioral data: for example, reaction times or perceptual accuracies in visual detection tasks are modulated at time scales scales of theta (∼4 Hz) and alpha (∼8 Hz) band activity [13-16]. In the case of hearing, neuroimaging studies have similarly shown that pre-stimulus delta (∼1 Hz) and theta activity (2-4 Hz) determine whether a sound is detected or influence how it is perceived [11, 12, 17-20]. As in vision, the influence of rhythmic activity manifests also in behavioral data [21, 22]. In general, the apparent influence of rhythmic neural activity on behavior has been linked to rhythmic modes of perception, which facilitate the amplification of specific, e.g. expected, stimuli and mediate the alignment of endogenous neural activity to the regularities of structured sounds such as speech [23, 24]. Indeed, the time scales of human perceptual sensitivity and the time scales at which rhythmic auditory activity shapes perception seem to be well matched [25, 26].

While it remains unclear whether truly spontaneous brain activity affects auditory perception [27, 28], it is clear that once the auditory system is driven by sounds rhythmic activity becomes engaged and shapes perception [12, 28-30]. Still, most studies linking neural activity and auditory percepts have relied on brief acoustic targets. Yet, a key prediction from models postulating a rhythmic mode of perception [23, 28], and from models linking cortical delta activity with sensory gain [3, 10], is that this rhythmic activity should also shape how sensory information is integrated over time: the perceptual weighting of temporally dispersed acoustic information should be modulated at precisely those time scales at which neural activity has been shown to shape the detection or perception of brief and isolated sounds.

We here tested this prediction directly at the level of behavior. That is, we asked whether the use of acoustic information available in a continuous and extended (1s or longer) stimulus is structured rhythmically at the delta/theta band time scale. To this end we employed an acoustic variant of the frequently used visual random dot motion stimulus [31]. In our study, participants judged the overall direction of change in the perceived frequency content of soundscapes composed of a dense sequence of random tones, a specific fraction of which systematically in- or de-creased in pitch (Fig. 1A). The level of sensory evidence available in each trial about the direction of frequency change was sampled independently in epochs of between 90ms to 180ms (in different versions of the task) allowing us to quantify the influence of the moment by moment varying acoustic evidence on participant’s judgements (Fig. 1B). Across four versions of this experiment we found consistent evidence that the perceptual sampling of temporally extended sounds is structured by processes operating at the time scale of around 1-2 Hz.

**Figure 1:**
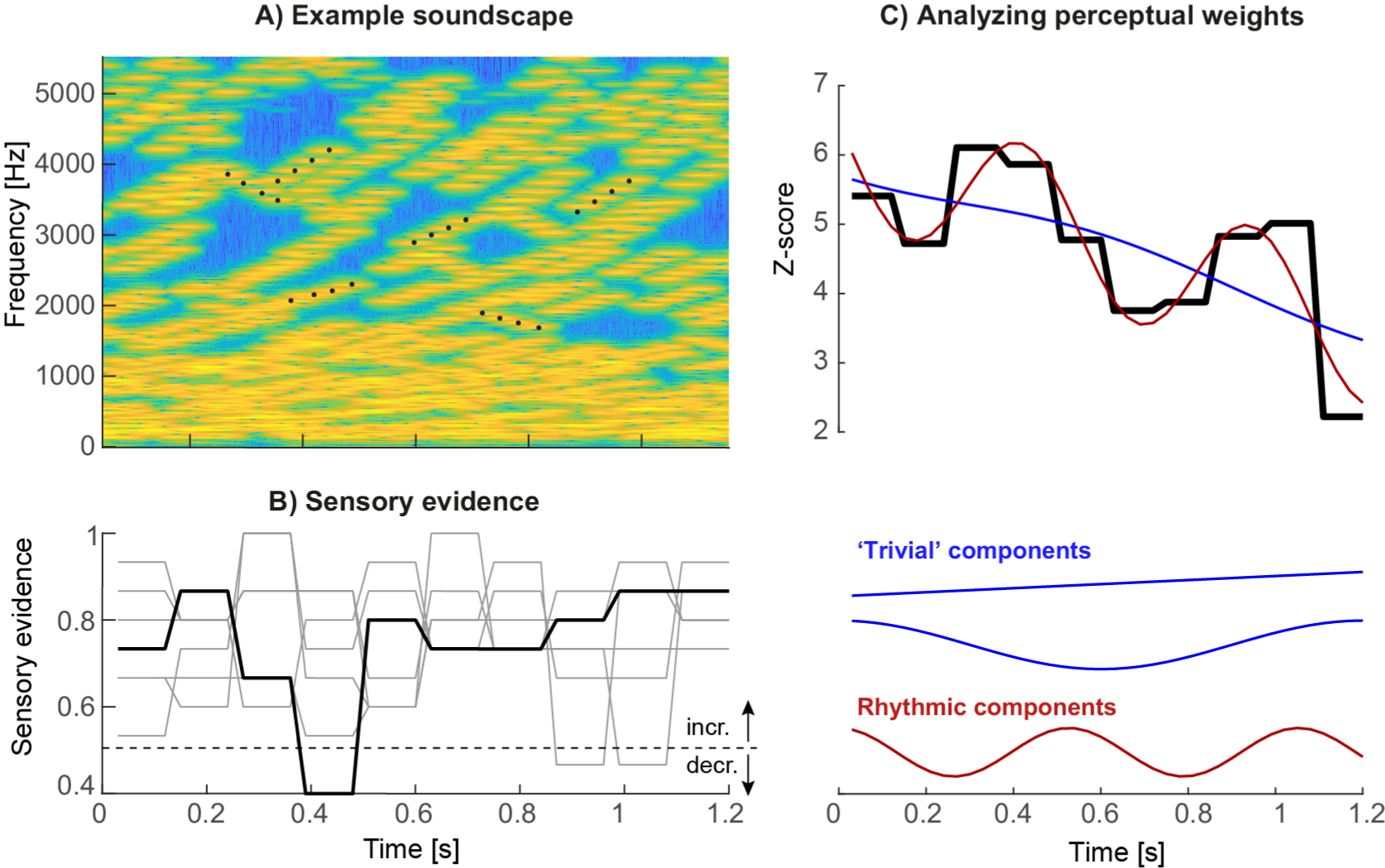
Acoustic stimuli and analysis. **A)** Stimuli consisted of ‘soundscapes’ consisting of 30 four-tone sequences either in- or decreasing in frequency (example sequences are marked by black dots). The fraction of sequences moving in the same direction changed randomly across trials and between ‘epochs’ of a specific duration, which varied between experiments. **B)** Each trial was characterized by the effective evidence for the soundscape to in- or de-crease, with the evidence being independent between epochs and trials, and varying around a participant-specific threshold. The black line presents the evidence for the soundscape shown in panel A, the gray lines the evidence for other trials, all with increasing frequency. Evidence levels of one correspond to a fully coherent increasing soundscape, levels of zero to a fully coherent decreasing soundscape. **C)** The trial-averaged single participant perceptual weights (average sensory evidence for trials where participants responded with ‘up’ or ‘down’, combined after correcting the sign of down responses) were analyzed using regression. These models distinguished trivial temporal structure arising from linear trends or U/V shaped profiles locked to stimulus duration from rhythmic structure at faster time scales. The graph displays the perceptual weight of one example participant together with the best-fitting trivial and rhythmic contributions.

## Methods

A total of 79 volunteers (age: 19 to 28 years, 67% female) participated in this study, following written informed consent and briefing about the purpose of the study. They received either monetary compensation or course credits. Seven individuals participated in two of the four experiments. All had self-reported normal hearing. The study was conducted in accordance with the Declaration of Helsinki and was approved by the local ethics committee of Bielefeld University. The required sample size per experiment was set a priori to at least n=20 based on recommendations for behavioral studies [32]. For two of the experiments an n of 23 was collected as data collection proceeded partly in parallel.

### Acoustic stimuli

Stimuli were presented via headphones (Sennheiser HD200 Pro) at an average intensity of 65 dB root mean square level (calibrated using a Model 2250 sound level meter; Bruel&Kjær, Denmark) while participants were seated in an acoustically shielded room. Stimulus presentation was controlled from Matlab (Mathworks) using routines from the Psychophysics toolbox [33].

The acoustic stimuli (‘soundscapes’) consisted of 30 simultaneous sequences of pure tones (each tone had 30ms duration, 6ms on/off cosine ramps) that either increased or decreased in frequency and each sequence lasted four tones (see Fig. 1A for a spectro-temporal representation). The initial frequency of each sequence was drawn independently (uniform between 128 Hz and 16’384 Hz), and each sequence in/decreased in steps of 20 cents. To construct a specific soundscape, each tone sequence started at a random position within the four-tone sequence, so that the start time of each sequence was independent of that of all others. Once a sequence reached the 4th tone, it was replaced by a new sequence starting at a random frequency. The impression of an overall in- or decrease in frequency over time was manipulated by changing the fraction of sequences that in- or decreased. This fraction (‘motion evidence’), was coded as between 0 and 1, with 0.5 indicating half the sequences as in- and the other half as decreasing, and 1 indicating that all sequences increased (see Fig. 1B for example traces). Each trial was hence characterized by the intended direction of change (in- or decrease) and the respective level of motion evidence at which this direction was expressed (i.e. the deviation from the ambiguous evidence of 0.5).

To quantify the perceptual use of the acoustic information at different time points by individual participants, this motion evidence was manipulated two folds. First, for each participant we determined (in an initial block) the participant specific perceptual threshold using three interleaved 1-up 2-down staircases. Second, we manipulated the motion evidence between trials, and over time within a trial, around this subject-specific threshold. To this end, we sub-divided each soundscape into ‘epochs’ and randomly and independently sampled the motion evidence from a Gaussian distribution around the participant-specific threshold (SD of 0.15 or 0.25, depending on the experiment). The duration of these epochs varied between experiments from 90ms to 180ms (see Fig. 1B for examples; cf. Table 1). Practically, for a given trial, it was first determined whether the soundscape should in- or decrease. Then, the epoch-specific levels of motion evidence were drawn and then the sequences of individual tones were generated as described above. The total duration of each soundscape varied between 1200ms and 1800ms (Table 1). Each experiment consisted of 800 trials per participant. For technical reasons, in some of the earlier experiments it had not been enforced that participants could respond only after the end of the soundscape, leading to premature responses. We hence imposed a minimal number of 750 valid trials for a participant to be included. Based on this criterion we analyzed the data of n=23 participants for experiments 1 and 2, and n=20 each for experiments 3 and 4 (Table 1).

**Table 1:**
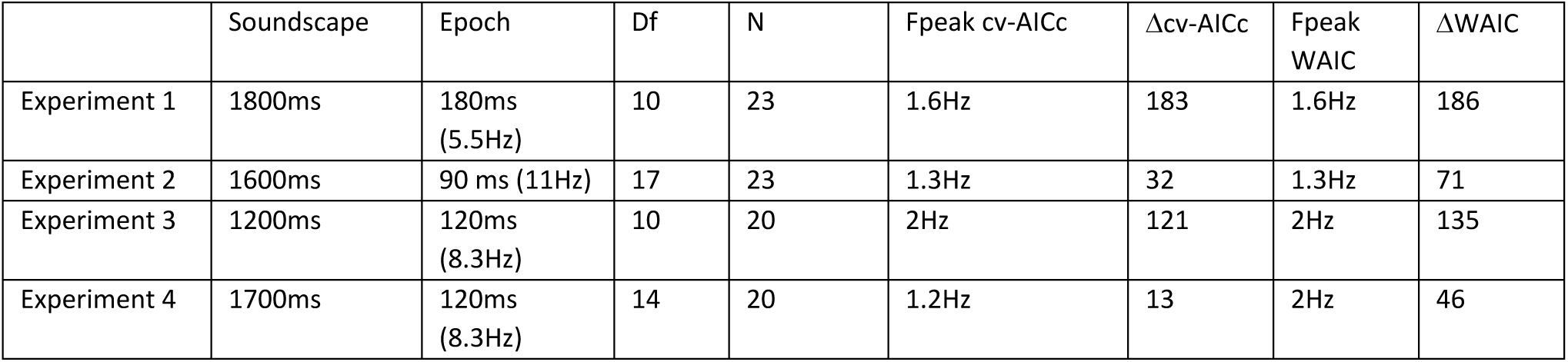
Parameters and results for each experiment, incl. the duration of sound scape and the ‘epochs’ over which the sensory evidence changed randomly, the number of epochs (sampling frequency) over which the perceptual weights were determined (Df), the number of participants (N), the best frequency determined by each model criterion (Fpeak) and the relative model criterion vs. the best trivial model.

### Response templates

In the domain of motion evidence each trial consists of a sequence of statistically independent samples of evidence for the direction of change in the soundscape. Hence, in this domain one can use the epoch by epoch evidence in a psychophysical reverse correlation procedure [34, 35]. To compute perceptual weights (also known as response templates) trials were split according to direction of change and participants’ responses. Then response-specific averages of the motion evidence (coded as deviation from the ambiguous evidence of 0.5) were computed and were converted to units of z-scores within each participant using bootstrapping; for this the actual weights were standardized relative to the distribution of 4000 sets of surrogate weights [36, 37]. These weights indicate how strongly the acoustic evidence at each moment influences the perceptual judgements, with zero indicating no influence and positive values indicating that the in- (de)creases in the stimulus were rated as in- (de)creasing by the participant (c.f. Fig. 2A). To visualize the spectral composition of these templates we computed their power spectrum after standardizing the overall signal power (Fig. 2B).

**Figure 2:**
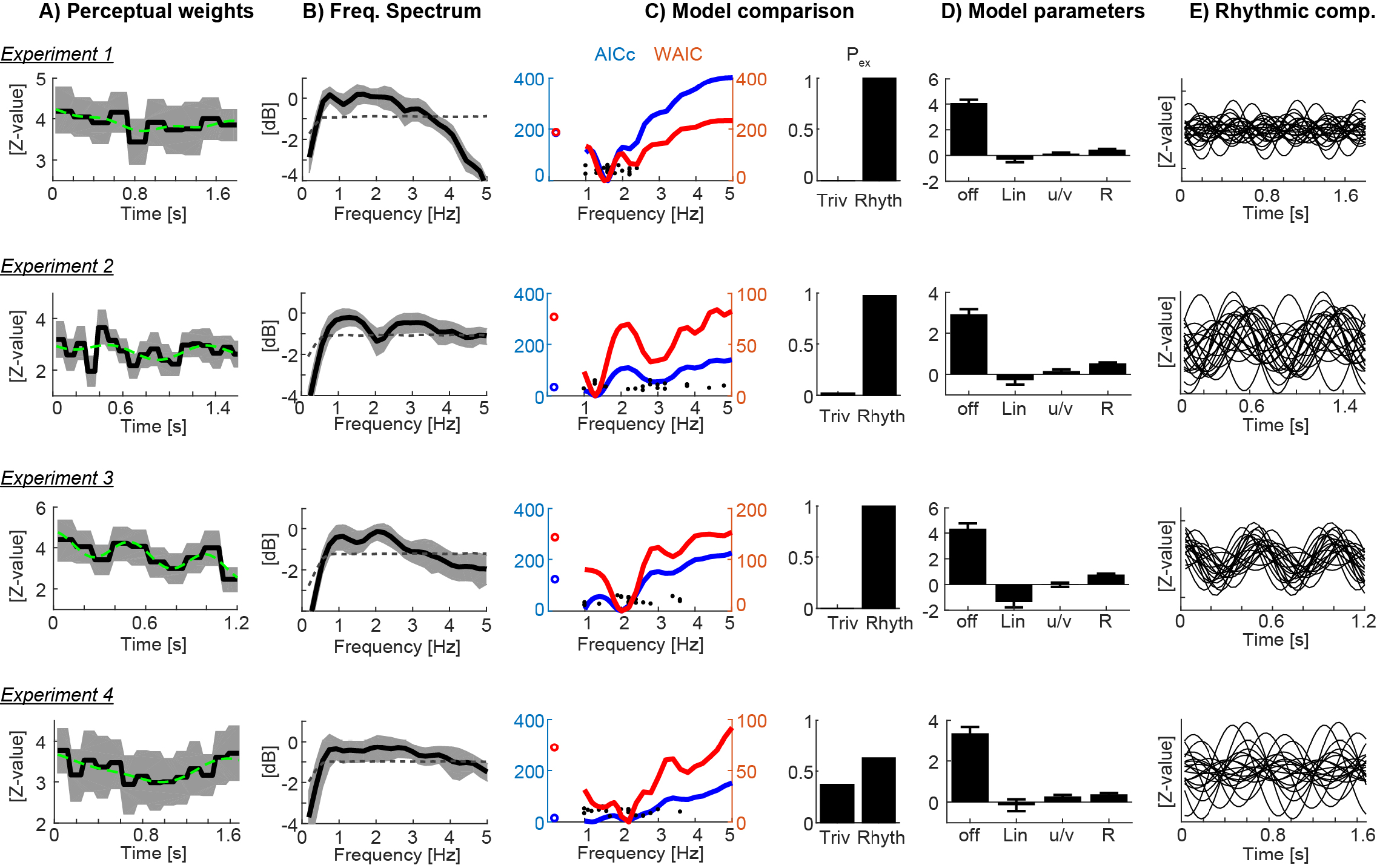
Results. **A)** Participant-averaged perceptual weights (black solid) with group-level two-sided 5% bootstrap confidence intervals (gray area) and the best-fitting model (green dashed). Units are in z-scores relative to a bootstrap baseline. **B)** Participant-averaged frequency spectrum of the perceptual weights (black solid), with two-sided 5% bootstrap confidence interval (gray area). The dashed line represents the average spectrum obtained after time-shuffling the weights. **C)** Model comparison results. The left curves show the group-level cv-AICc (blue) and WAIC (red) values for the best trivial model (open circles) and frequency dependent models. Individual participant’s best frequencies are denoted by solid black dots. The right bars show the exceedance probabilities of a comparison between the trivial model and the rhythmic model derived at the frequency yielding the lowest group-level cv-AICc. **D)** Model parameters of the trivial contributions (offset, linear slope, u/v shaped profile) and the rhythmic component (R, root-mean-squared amplitude of sine and cosine components). Bars and error-bars indicate the group-level mean and s.e.m. **E)** Rhythmic component of the best-fitting model for each participant.

### Analysis of rhythmic components in response templates

The main goal of the analysis was to determine whether the participant-specific weights were characterized by a significant rhythmic component (see Fig. 1C for example weights). To this end an analysis was implemented that selectively compared non-rhythmic structure, such as i) the overall offset, ii) a linear in- or de-crease over time, and iii) a U/V shaped profile time-locked to the stimulus duration (modelled as cos(2*pi*t*F_exp_), with F_exp_= 1 / stimulus duration). For each participant’s perceptual template, we first computed regression models comprising a collection of the non-rhythmic patterns (only i, i+ii, and i+ii+iii) and selected from these best fitting model based on the group-level AICc (see below). We then extended this ‘trivial’ model by a rhythmic contribution modelled at frequencies varying between 1 to 5Hz, comprising both sine- and cosine components of the same frequency. We then used formal model-comparison approach to determine, at the group-level, whether there was evidence for the rhythmic model to better explain the data than the trivial model. Comparison was based on the log-likelihoods computed for each participant’s data and regression model.

Two recommended approaches to model comparison were used [38, 39]. First, regression models were fit using a Monte-Carlo approach to compute the Watanabe-Akaike information criterion (WAIC), which captures the out-of-sample predictive power when penalizing each model [39]. This calculation was implemented using the Bayesian regression package for Matlab [40], using 10’000 samples, 10’000 burn-in samples and a thinning factor of 5. Second, regression models were compared using a cross-validated version of the corrected Akaike criterion (cv-AICc) [41]. Response templates were fit using half the data and the log-likelihood of each model was computed on the other half; the AICc was then averaged over 10 independent 2-fold CV runs. Group-level comparison was performed by computing the group-level WAIC and cv-AICc, and by computing the exceedance probabilities of each model based on −0.5*cv-AICc (implemented using VBA toolbox in Matlab [42]).

To determine whether the selection of a specific frequency-dependent model over the trivial model was indeed specific to the behavioral data, and was not induced by any other factor in the analysis such as the temporal binning (see e.g. [43] for pitfalls involved in testing reaction times for rhythmic patterns), we repeated the model comparison using randomized data. We randomly paired stimuli and responses and computed the probability of selecting the rhythmic model (at the group-level frequency determined using the original data) over the trivial model across 2000 instances of randomized data based on the cv-AICc.

## Results

Participants judged the perceived direction of frequency change in soundscapes constructed based on randomly varying task-relevant evidence. Across four variants of the experiment the stimulus duration ranged from 1200 to 1800 ms, while the time scale at which perpetual weights were sampled varied between 5.5 and 11 Hz (Table 1).

Figure 2A displays the group-averaged perceptual weights for each experiment. The weights were significant for all time points (based on the 5% percentile of a bootstrap distribution), in line with the task difficulty being set to be around each participant’s perceptual threshold. For each dataset, the group-level weights exhibited a rich structure, comprising linear trends and a U/V shaped profile tied to the duration of each soundscape. To determine the contribution of such ‘trivial’, i.e. non-rhythmic, contributions we fit three candidate models to each participant’s data. Group-level model comparison revealed that the model featuring all three trivial factors (offset, slope, U/V profile) better explained the data than a reduced model: the group-level ΔAICc of the full vs. the reduced model were 211.2, 4.6, 93.3, 117.2 for experiments 1 to 4 respectively, and the group-level exceedance probabilities of the full model were p_ex_=0.76, 0.49, 0.99, and 0.94. Only for dataset2 there was no clear evidence for any of the models to explain the data better than the others.

We then used the best trivial model (determined at the group-level, separately for each experiment) to quantify the extent to which the addition of a rhythmic contribution helped to better explain the perceptual weights. The prominence of temporal structure at the time scale between 1 and 3Hz is also highlighted by the frequency spectra in Fig. 2B. Formal model comparison between the best trivial model and the frequency dependent models revealed that the addition of a rhythmic component between 1.2 and 2Hz significantly improved the model fit (Fig. 2C). The time scales best explaining the perceptual data were 1.6Hz, 1.3Hz, 2Hz and 1.2Hz based on the cv-AICc for the four experiments respectively (Table 1). When using the WAIC we found the same frequencies, except for experiment 4 (here WAIC identified 2Hz as best frequency). Both the cv-AICc and the WAIC identified the rhythmic model (defined at the group-level AICc-based best frequency) as significantly better than the trivial model for each dataset: the Δcv-AICc values of best rhythmic over the trivial model were 183, 32, 121, 13 respectively, the ΔWAIC values were 186, 71, 135, and 46. To further substantiate this result we obtained group-level exceedance probabilities of the best rhythmic model in comparison to the trivial model: for three out of four experiments these clearly favored the rhythmic model: p_ex_= 1, 0.97, 1, 0.62 for experiment 1 to 4 (Fig. 2D).

Given that this apparent rhythmic structure may also emerge simply as byproduct of sub-sampling the behavioral sensitivity at a fixed time scale, we repeated the model fitting after shuffling behavioral responses across trials. We computed the probability that the model incorporating the best group-level frequency derived from the original data better explained the data than the trivial model in the shuffled data (based on the cv-AICc): these probabilities were small and revealed the actual effects as (close-to) significant: p=0.08, 0.076, 0.040, and 0.068 for experiments 1 to 4 respectively.

To visualize the best models, panel 2E displays the model parameters for the best-fitting rhythmic model, while panel 2F displays the rhythmic component for each individual participant.

Closer inspection of Fig. 2C shows that the WAIC reveals two local minima for several of the experiments: besides the overall best model at frequencies between 1.2 and 2Hz, also rhythmic components at frequencies between 2 and 4Hz better explain the actual data than the trivial model. The precise frequency of this second component varied between experiments (experiment 1: 2.4 Hz, ΔWAIC = 123 vs. trivial model; experiment 2: 2.8 Hz ΔWAIC = 43; experiment 3: 3.4Hz ΔWAIC = 36; experiment 4: 3.8Hz ΔWAIC = 24). This observation suggests that effectively multiple rhythmic components may underlie auditory perception.

## Discussion

We investigated whether the relation between the sensory evidence contained in temporally extended soundscapes and participant’s judgements is governed by rhythmic components, as predicted by theories of rhythmic models of listening and studies linking delta/theta band neural activity with perception. The four experiments differed in the overall stimulus duration (Table 1; 1200 ms to 1800 ms) and the time scale at which perceptual weights were sampled (5.5 to 11 Hz). Despite these variations in the experimental paradigm we found converging evidence that the perceptual sensitivity profiles contain significant rhythmic structure at the time scales between 1.2 and 2 Hz. That the rhythmic models indeed better explain the perceptual use of acoustic information than a trivial model only containing linear and U/V shaped trends is supported by the use of two criteria for formal model comparison and the comparison of the original and shuffled data.

The perceptual weights featured pronounced non-rhythmic temporal structure, such as linear trends (e.g. experiment 3, Fig. 2A) or a U-shaped profile emphasizing early and late stimulus components (e.g. experiment 4). Such stimulus-locked temporal sensitivity is frequently observed in perceptual decision making paradigms and in part may reflect the participant-specific strategies for analyzing the sensory environment, temporal leakage in decision processes, or the urgency to respond [44, 45]. Importantly, our results show that this sensitivity profile is augmented by a more rapidly changing temporal structure that emerges at precisely those time scales deemed relevant for auditory perceptual sensitivity by neuroimaging studies.

Consistently across the four experiments, the best rhythmic models featured a perceptual sensitivity that was modulated with a frequency between 1.2 and 2Hz. Previous work has shown that auditory cortical delta band activity is tied to changes in the network state related to an overall rhythmic fluctuation in neural background activity, visible both in spontaneous and acoustically driven states [3, 4]. In particular strong engagement of auditory delta band activity has been implied in acoustic filtering of attended information and the task-relevant engagement of auditory networks [29, 46, 47] and plays a central role in theories of rhythmic modes of listening [23, 28]. While electrophysiological studies reporting behaviorally-relevant rhythmic patterns of brain activity often identified frequencies in the theta band as important [11, 12, 17, 48] some of these have identified multiple mechanisms operating at different time scales, including the delta band between 1 and 2Hz [12, 19, 49]. Our results corroborate the behavioral relevance of neural mechanisms operating in the delta band for auditory perception and provide evidence for the potential existence of distinct and possibly multiplexed rhythmic mechanisms.

Still, there are a few caveats to note. First, while the converging evidence across the four experiments is very convincing, for each individual experiment the statistical likelihood of the rhythmic model to better explain the data than the trivial model in comparison to randomized data was only marginally significant. One possibility is that the estimated perceptual weights are still noisy and more trials per participant would be required to obtain fully reliable estimates. Second, it could be that the preferred perceptual sampling frequency differs across participants, precluding a reliable estimate of a common group-level model. Indeed, the single-participant data reveal a considerable variability in their best-frequency (c.f. Fig. 2C). However, without the assumption of a fixed group-level frequency it becomes difficult to determine whether a frequency dependent model fits the data significantly better than a null model. Third, the presented analysis implicitly assumes that the participants made use of the full available acoustic information and used all tone sequences equally for their judgements. The perceptual reverse correlation procedure was implemented in the domain of the overall motion-evidence, based on which the different tone sequences were randomly assembled. Performing a reverse correlation in the full time-frequency domain would likely require much higher trial numbers as the degrees of freedom for the perceptual weights would increase considerably. As a result, participant specific biases towards particularly low or high sound frequencies may have reduced the power of the present analysis. Considering the degrees of freedom of the analysis, that is the number of effective weights per perceptual template, Table 1 reveals that the two experiments with the lowest number of free parameters were those yielding the larges evidence in favor of a rhythmic model, regardless of the total duration of the soundscape. This observation fits with the possibility that when sampling perceptual weights at finer temporal resolution or over additional dimensions, such as sound frequency, more trials would be required to obtain reliable estimates.

Also, the present results leave it unclear whether the rhythmic process(es) operate at the level of sensory encoding or decision making. When combined with fixed-duration stimuli, psychophysical reverse correlation cannot dissociate sensory from decision processes [44]. While electrophysiological studies have directly demonstrated the relevance of auditory cortical theta band activity for neural sound encoding [3, 47] and perception [29], neuroimaging studies have shown that rhythmic brain activity may affect both the encoding of sensory information at shorter latencies and decision processes emerging later in frontal regions [17, 49]. Work on visual decision making has also demonstrated the relevance of delta band activity for the accumulation of sensory evidence over time [6]. Hence it could be that the rhythmic patterns revealed here either reflect a change in the quality of the encoding of sensory evidence at each moment in time, which then results in a differential contribution to the participant’s judgement, or a direct change in the weight assigned during the accumulation of evidence for choice [6, 50]. More work is required to better understand the interplay of rhythmic processes related to sensory encoding and of those related to the actual decision process.

In sum, theories of rhythmic modes of listening and neurophysiological data linking network activity to single neuron encoding predict that rhythmic activity shapes how acoustic information is combined over time to judge extended soundscapes. The present study proposes one approach to test this and provides converging evidence in support of this prediction. Future work can capitalize on this approach to directly link electrophysiological signatures of rhythmic activity to the perceptual combination of acoustic information over time.

## Acknowledgements

I am thankful for Sophia Schreiber and a number of students for collecting the data.

## Data and code availability

The behavioral data and the required Matlab code for producing the stimuli, the analysis and figures are available from http://www.uni-bielefeld.de/biologie/cns/resources.html (section The rhythmic sampling of auditory scenes).

